# Automated, high-throughput kinematic analysis of skilled reaching in rodent models of neurological injury

**DOI:** 10.64898/2026.11.13.766238

**Authors:** Logan S. Friedrich, Esmeralda Mendoza, J. Russell Huie, Jose Rosas, Zimei Wang, Sunil Mathew, Colin Schultz, Neil Chhikara, Nick Samburg, Riley Specht, Isabel Antolak, Pantelis Tsoulfas, Adam R. Ferguson, Murray G. Blackmore

## Abstract

Neurological dysfunction can diminish quality of life by disrupting the movement capacity required to accomplish tasks. Kinematic studies guide treatment development by revealing subtle relationships between patterns of movement and neurological dysfunction. However, these studies produce a large quantity of opaque, multifaceted, and often highly correlated data. Here we present a novel and efficient approach to collecting, analyzing, and interpreting kinematic data with high throughput. We automated a single pellet reaching task in mice and created a video analysis pipeline that scores task success and extracts kinematic features from reaches. To analyze the resulting data, principal component analysis was first applied to group covarying measurements into a few overarching dimensions of behavior. Regularized regression was then applied separately to select reach qualities that best predicted task success. Applied to four nervous system injury models, this approach revealed differences in the effects of injury type and rehabilitative intervention on movement patterns that were not apparent from task success alone.

## INTRODUCTION

Turning large, complex datasets into specific targets for therapy and clear measures of recovery is a central challenge in neural repair research. Doing so requires meeting at least three main objectives. First, data must be captured on a sufficient scale. When the area of inquiry involves large inter-subject variability, which is the case in reparative neuroscience (Button et al., 2013; Fouad et al., 2021; Watzlawick et al., 2019), achieving scale entails both deep capture of information from each subject and workflows with sufficient throughput for adequate power. Second, methods are needed to identify relationships in high-dimensional data and to simplify presentation and manipulation. This objective can be met with dimensionality-reducing approaches like principal component analysis (PCA) (Cunningham & Yu, 2014; Ferguson et al., 2013). However, although valuable for downstream applications like enhanced prediction and classification, the resulting data constructs are highly abstract and may not lead to concrete hypotheses or specific optimization steps (Jollife & Cadima, 2016; Ma & Dai, 2011). This raises a third objective, which is to link specific data elements to desired outcomes, identifying tangible targets for follow-up study and providing a rational framework to prioritize subsequent attention.

Here we address each objective in a study of forelimb impairment after various forms of nervous system damage and subsequent rehabilitation. To address the need for data at scale, semiautomated administration of the single pellet retrieval task (SPRT) and automated kinematic tracking in digital recordings enabled profiling of more than 150,000 reach attempts from 62 animals across thousands of pre-injury, post-injury, and post-rehabilitation testing sessions. To identify relationships and simplify the resulting high-dimensional data, we employed principal component analysis (PCA), identifying groups of co-varying elements related to motion characteristics and, in a separate component, indicators of task engagement. Finally, for more targeted insights, we applied least absolute shrinkage and selection operator (LASSO) regression to reveal specific kinematic features most strongly associated with retrieval outcomes, enabling specific tracking of key predictors of success after injury and rehabilitation (Quinlan et al., 2015; Tibshirani, 1996). This integrated collection of hardware, algorithmic data extraction, and complementary analysis pipelines offers a A flexible approach to rapidly generate rich behavioral data, identify the reach features most closely tied to task success, and test whether interventions restore them.

## MATERIALS AND METHODS

### Animals

All procedures were approved by the local Institutional Animal Care and Use Committee (IACUC) and were conducted in accordance with the NIH guidelines for the care and use of laboratory animals. In total, these experiments used 120 mice of both sexes that were 8-10 weeks of age when training began, of which 36 were removed due to mortality or failure to acquire the behavioral task, leaving 84. Mice were obtained by homozygous breeding from Emx1-Cre founders (C57/Bl6 background) obtained from Jackson Labs (#005628, RRID:IMSR_JAX:005628). No cre-activated transgenes were supplied in these experiments and expression of cre, which is confined to forebrain cells in this line, is not known to affect cellular physiology or animal behavior, leaving animals effectively wildtype C57/Bl6 (H. Guo et al., 2000; Wang et al., 2026). Mice were group housed on a 12-hour light/dark cycle with light on at 6:00 AM and were tested during the light phase. To motivate reaching, access to food was restricted to a ∼2.5 hour feeding window, with feeding opportunity adjusted to maintain between 85 and 90% of starting body weight. Restrictions were lifted following the final behavioral session of each week and resumed the day prior to testing. Mice were returned to ad libitum feeding for at least 3 days before surgery and at least 7 days after surgery. All quantification was performed by personnel blinded to animal identity.

### Animal injuries

All injuries were performed under deep general anesthesia induced by a ketamine/xylazine mixture (100 mg/kg; 10 mg/kg i.p., respectively). Immediately following surgery, mice received a subcutaneous injection of meloxicam (5 mg/kg) as post-operative analgesia. Post-surgical monitoring occurred for at least the following 7 days and additional meloxicam was provided for the following 3 days. Return to behavioral testing occurred no sooner than 10 days after any surgical procedure was performed.

Dorsal hemisection and contusion injuries shared a common dorsal cervical surgical approach. The hair over the dorsal neck was removed and the surgical site cleaned with povidone-iodine solution and ethanol wipes. To access the spinal column, an incision was made and musculature laterally retracted to expose the C5 and adjacent vertebrae. Mice were mounted in a custom spine stabilizer (Wang et al., 2025; Wang, Maunze, et al., 2018) and residual musculature was cleared enabling identification of C5. To expose the cord and enable injury, a C5 laminectomy was performed with Noyes spring scissors. Dorsolateral hemisection was performed at C5 on the right dorsolateral aspect of the cord using a vibraknife set to a depth of 0.8 mm while avoiding visible vasculature (Wang et al., 2015, 2017; Wang, Mehra, et al., 2018). Contusion was performed with a force-controlled impact delivered to C5 by an Infinite Horizon Impactor (IH-0400) aligned slightly right of midline. Force was set to deliver 60 or 70 kdyn depending on group with a 0 second dwell time. Unilateral pyramidotomy was performed following the procedures of (Wang et al., 2015). Using a ventral approach, the medullary pyramids were exposed by removal of the ventrocaudal region of the occipital bone with delicate rongeurs. The dura was punctured, and the left pyramid was cut completely using a micro feather scalpel.

### Automated single pellet retrieval apparatus (ASPA)

Mice performed a single pellet retrieval task (SPRT) from within custom plexiglass enclosures based on the Farr-Whishaw design, which has a 5 mm slit in the front-right corner of the box (Farr & Whishaw, 2002). A wall adjacent to the slit permitted reaching for the pellet with the right forelimb while suppressing the ability to reach with the left forelimb. A motorized conveyor carried a 3D-printed tray bearing 20 pellet-containing segments, each containing a millet seed atop a divot and isolated from neighboring areas by walls. A NEMA-17 bipolar stepper motor (17HS19-2004S1, Stepper Online) was controlled by an Arduino microcontroller equipped with a custom shield and driven by a stepper motor driver to advance the tray to present the next pellet every 30 seconds, presenting each pellet in turn in alignment with the slit. A piezo buzzer signaled each advance, and the operator made minor adjustments to improve pellet alignment with the slit as needed. Trays were double-sided, holding 20 pellets per side. After the final position, the belt paused and the operator reversed the tray to the second face before the belt returned to its origin. Because four enclosures were placed along each belt and four such belts were stacked on a shelving rack, a single operator tested as many as 16 animals in parallel. A generic webcam was uniformly mounted above each enclosure using a custom 3D-printed mount. Video feeds were assembled into a live collage view with each capture dimension made uniform. For each tray side, the live collage was recorded by screen capture at 1920 × 1080 pixels and 60 frames per second, so that each animal’s view occupied 480 × 540 pixels. These collages would later be cropped into 8 single mouse videos for offline analysis. Complete details about the device, including construction resources, are available through the GitHub repository associated with this project: https://github.com/RegenerationLab/SPRT.

### Tray geometry

Pellets were presented on 3D-printed trays of 4 varieties, each presenting alternate pellet presentations and difficulties. All presentations spaced divots (on which millet seed pellets were placed) 1 cm apart and isolated the adjacent presentation areas with walls. Between types, trays differed in divot closeness to the cage slit as well as features surrounding the divot which progressively increased difficulty of retrieval and either allowed or prohibited additional attempts once a pellet was displaced from its divot. “Easy” trays featured a closer back wall, closer divot location, and a flattened surrounding area that enabled reattempts. “Flat” trays moved the wall and divot location further away from the slit but otherwise resembled easy trays. “Recessed” trays resembled flat trays except the divot was centered within a 3 mm wide by 1 mm deep cylindrical depression. “Pillar” trays resembled flat trays except the entire area surrounding the divot was depressed creating a pillar on which the pellet rested. A wall immediately adjacent to the slit opening and rising to the level of the pillar prevented access to the collection area and thereby prohibited retrieval if the pellet dropped. Thus, scooping and dragging solutions were prevented and success on the pillar tray required pellet grasping for retrieval to succeed. The pillar geometry was selected in a pilot experiment in which a separate group of 22 female mice was tested on a single tray type each (flat, n = 7; recessed, n = 8; pillar, n = 7) before unilateral pyramidotomy and weekly for eight weeks afterward, without rehabilitation (Supp. Fig. 1; Table 1). Injury, pellet scoring and histological verification were performed as described within the main experiment. Trays were loaded in a single rapid step with the aid of tray-specific loading shields. Once placed over the tray, the assembly was dipped into a container of millet seeds and excess pellets were shaken off. Upon removal of the shield, one pellet remained in each divot, thereby simultaneously loading all 40 positions of each tray. Full designs of all tray types are available on GitHub: https://github.com/RegenerationLab/SPRT.

### Behavioral training and testing

Behavioral shaping occurred to encourage reaching through the slit to retrieve pellets. Initial training involved placing the mice into boxes with food pellets presented at the slit via a 3D-printed inclined vat loaded with approximately 3 g of pellets. The slope of the vat ensured constant pellet access throughout a 1-hour administration window occurring across 3 days or until mean retrieval of 1 g of pellets. All behavioral testing following this timepoint was recorded on video for later analysis. Animals then transitioned to pillar trays for further training, four pillar-tray sessions (80 pellets total) per day, until they reached a performance plateau. Baseline / pre-injury performance was calculated as the average of the final three days of training. According to the pellet’s position on the tray at the end of administration, task outcomes were manually scored as either retrieved if missing, displaced if within the tray segment while not on the divot/pillar, or missed if present on the pillar. Following injury, mice received 4 weeks of post-injury testing on 40 pellets presented on pillars, once per week. This was chosen to avoid inadvertent rehabilitation and practice effects. Following post-injury testing, mice received activity-based rehabilitation in which progressively more demanding trays were presented over the course of 2 weeks, each day consisting of 2 sides of the same tray type: easy trays for 5 days and flat trays for 3 days. During the following 2 days, mice were presented with pillar trays, and this period was used to assess rehabilitation-mediated recovery.

### Automated reach analysis

To study motions used while performing SPRT, videos were recorded and subsequently analyzed with a reach kinematics pipeline. Videos were renamed to a specific format according to the date, the location of each mouse within each box and within the video, followed by the tray type (denoted by the first letter of each tray type) and tray repetition for that day. A cropping script turned these collages into 8 single-animal videos by pixel coordinates using FFmpeg and renamed them according to their location (using the mouse in the upper right corner of the collage video as an example: 20251201_D04_P3.mp4). These single-animal videos were then processed with a DeepLabCut neural network that identified various points on the mouse and in its environment: the distal tip of the most extended digit which we called the location of the right forepaw, a 3D-printed reference point that was mounted to each box in fixed position, the pellet in the scoring segment directly in front of the slit, the pillar when and only when the same pellet was not on that pillar, the tip of the mouse’s nose, the point on the ears most distal to the nose, the other paws, and the tail base. The model was a ResNet-50 network trained on about 10,000 hand-labeled frames selected at random from the video corpus. Tracking accuracy was evaluated on hand-labeled frames that were withheld from training. Of the labeled frames, 95% were used to train the network and 5% were reserved for testing. After 1,030,000 training iterations, the mean distance between predicted and hand-labeled positions was 1.92 pixels on the training frames and 3.43 pixels on the withheld test frames. The same network was applied to every video in the study. The resulting output was a spreadsheet of frames within the video, and the Cartesian coordinates most likely to contain each tracked object as well as the likelihood score. These outputs were parsed with a custom script package that identified each reach. A reach was a run of frames in which the nose was within the slit region and the right forepaw lay at least 8 pixels beyond the nose along the reach axis, the image axis running from the enclosure through the slit toward the pellet. It began at the first such frame and ended at the last one before a gap of two or more frames, or when the paw stopped moving. It also detected changes in the pellet’s location or its disappearance. Together, these processes identified which reach caused which pellet outcome. The location of the paw throughout the duration of each reach was then used to calculate a series of kinematic features: reach length (peak paw excursion along the reach axis), breadth (lateral excursion), path length (the summed frame-to-frame travel of the paw), the area enclosed by the paw path, speed, and duration. Kinematic measures are reported in image units (pixels and frames) of the 480 × 540 pixel single-animal videos. The reference point served to locate the slit in each video and was not used to convert distances to physical units. Attention paid to the task was also tracked as the percentage of the video’s duration during which the nose was within the slit. In the analysis, a tracked point was used only when its likelihood exceeded the criterion set for that measure (0.7 for the right forepaw in reach detection, 0.9 for the nose in the attention score, and 0.95 for the position of the pellet); points below the criterion were excluded and were not interpolated. Across 62 mice, the algorithm identified 159,343 reaches. Of these, 20,384 were made when no pellet was present at the presentation position, most often (84%) because an earlier reach had already retrieved or displaced it; these were assigned no outcome, leaving 138,959 scored reaches. Automated outcome counts were compared with manual tray scores in the 3,814 sessions for which both were available. Per-session counts were correlated for contacted pellets (r = 0.89; manual mean 9.9, automated mean 10.0), retrieved pellets (r = 0.70; 3.5 versus 3.2) and displaced pellets (r = 0.76; 6.4 versus 6.9). Models, training data, and analysis scripts may be accessed through the GitHub repository associated with this project: https://github.com/RegenerationLab/SPRT. Together, this pipeline yielded, for each session, an automated measure of attention alongside the outcome and kinematics of every reach, without the need for manual frame-by-frame scoring.

### Histological verification of injury

Mice were euthanized and fixed by transcardial perfusion with 4% paraformaldehyde (Electron Microscopy Sciences, 15710) prepared in phosphate-buffered saline (PBS; Sigma, P4474). The cervical spinal cord was dissected, kept overnight (16 hours) in the same fixative, and rinsed in 1X PBS before brain, medulla and cervical spinal cord were finely dissected. Tissue was then processed according to injury type.

Pyramidotomy was assessed for completeness by PKCγ immunolabeling. Cervical spinal cords were embedded in 6% gelatin and cut into 100 µm transverse sections with a vibrating-blade microtome (Leica VT1200). Blocking solution contained 10% normal goat serum (NGS; Gibco, 16210064) and 3% Triton X-100 (Sigma, 9002-93-1), and both antibodies were diluted in 3% NGS with 0.4% Triton X-100. The primary antibody was a rabbit anti-PKCγ (Abcam, ab71558, RRID:AB_1281066; 1:500). It was followed by a goat anti-rabbit secondary antibody conjugated to Alexa Fluor 647 (IgG H+L, cross-adsorbed; Invitrogen, A-21244; 1:500), applied at room temperature for two hours. Sections were mounted on slides and imaged at 20X on a Nikon Ti2-E inverted microscope fitted with a Nikon A1R+ laser scanning confocal system. To quantify PKCγ signal, the left and right dorsal columns were outlined by hand in the 20X images and the average pixel intensity inside each outline was measured. Intensity in the right (injured) column is reported as a percentage of intensity in the left (uninjured) column. Injuries were considered complete when intensity was reduced by at least 90% on the injured side; all surviving animals met this criterion.

Dorsal hemisection was verified by immunolabeling for glial fibrillary acidic protein (GFAP). Cervical spinal cords were embedded and sectioned as above, except that 100 µm sagittal sections were cut through the injury site. Sections were left overnight in a primary antibody against GFAP (Dako, Z0334, RRID:AB_10013482; 1:500), rinsed, and placed for 2 hours in an Alexa Fluor-conjugated secondary antibody (Thermo Fisher Scientific, R37117; 1:500). Images of the lesion were collected on the same confocal system.

Contusion was assessed by lesion volume in optically cleared tissue. Spinal cords were cleared with a modified 3DISCO protocol (tetrahydrofuran series followed by benzyl alcohol/benzyl benzoate) as described previously (Wang et al., 2022). Cleared tissue at the injury site was imaged on an Andor Dragonfly 202 spinning-disk confocal system mounted on a Leica DMi8 inverted microscope, using an HC PL APO 10×/0.45 objective. Tissue autofluorescence was excited with the 561 nm laser of an Andor Integrated Laser Engine at 15% power, collected through a 594/43 nm emission filter, and captured on an Andor Sona 4.2B-6 sCMOS camera with a 100 ms exposure. Z-stacks were acquired at a voxel size of 0.603 × 0.603 × 1.61-3 μm (x × y × z) and covered the full rostro-caudal extent of the lesion. Lesion volume was measured in three dimensions using Imaris version 10.2 (Oxford Instruments, Bitplane). Lesion boundaries were defined using tissue autofluorescence in the red channel. Injured tissue differed clearly from adjacent intact tissue: it showed lower signal intensity and an irregular, heterogeneous texture, whereas intact tissue had uniform signal and regular structure. Because automated intensity thresholding could not reliably separate the two, the lesion boundary was traced manually. Using the manual contour tool of the Imaris Surfaces module, the border between injured and intact tissue was drawn on every 4th to 5th optical section in the horizontal plane, across the full rostro-caudal extent of the lesion. Imaris then interpolated the contours into a closed three-dimensional surface, and lesion volume (μm^3^) was taken from the Surfaces statistics.

### Statistical analyses

Behavioral and kinematic outcomes were analyzed with linear mixed models. Examined outcomes included: retrieval, contact (retrieved plus displaced), attention, and each per-reach kinematic feature together with its derived principal components. For each outcome we fit a model with injury group, time, and their interaction as fixed effects and animal as a random effect, the random term accounting for repeated measures across timepoints. Effects were reported with eta-squared effect sizes, and post-hoc contrasts were computed from estimated marginal means. Pairwise comparisons among injury groups within a timepoint used Tukey adjustment; planned temporal contrasts, and comparisons of those changes between groups, used Holm adjustment. Within-group estimates are expressed as the later timepoint minus the earlier timepoint, so that negative values denote a decrease. Because retrieval approached a floor in the more severely injured groups after injury, we additionally verified these effects with a non-parametric approach. To identify which reach features strongly predicted retrieval, we applied LASSO feature selection (IBM SPSS v31.0). The LASSO penalty parameter (λ) was evaluated over a prespecified range from 0.00 to 1.00 in increments of 0.02. Model prediction error was estimated using 10-fold cross-validation. To favor parsimony while retaining comparable predictive performance, the final model was selected using the one-standard-error criterion, which chooses the more parsimonious model whose cross-validated prediction error is within one standard error of the minimum. We also used principal component analysis to reduce the set of correlated reach features to a smaller number of components for modeling. Animals that failed to acquire the task, defined as baseline (post-training) retrieval below 15%, were excluded before analysis, yielding a final sample of: pyramidotomy (n = 12), dorsal hemisection (n = 27), 60-kdyn contusion (n = 14), and 70-kdyn contusion (n = 9).

A non-linear principal component analysis (NL-PCA) was conducted using the syndRomics package to identify latent patterns while reducing the dimensionality of the dataset. Sixteen variables were used: video retrieved, video displaced, video contacted, total swipes AI, swipe speed, swipe length, swipe duration frames, swipe breadth, swipe area, path over frames, path length, manual retrieved, manual displaced, manual contacted, contacted match, and attention AI. Data were organized in long format with each row representing one animal at a single experimental session. To select the optimal number of principal components (PCs), a combination of methods was used, including the Kaiser criterion (eigenvalues greater than one), the elbow method that observes the asymptote in the scree plot, and bootstrapped permutation testing (Torres-Espín et al., 2021). A loading threshold of |0.45| was used to identify variables contributing to each PC by the directionality of individual variables and how they move along the PC for a PC phenotype label. Additionally, each animal session observation was assigned a PC score for each retained PC.

A two-way repeated measures analysis of variance (RM-ANOVA) was used to evaluate significant differences across PC phenotypes by injury group and time. Significant effects were reported using p-values, effect sizes were reported with an eta-squared, and post-hoc comparisons were conducted using estimated marginal means (EMMs), with Holm adjustment for differences among injury groups across time. All statistical analyses were performed in R version 4.5.2. Lesion volumes of the two contusion groups were compared with Welch’s two-sided t-test, which does not assume equal variances, with a Mann-Whitney U test as a non-parametric check; unlike the analyses above, these two tests were performed in Python (SciPy).

## RESULTS

### Semi-automated administration of the single pellet retrieval task

To increase throughput in the single pellet retrieval task (SPRT), a widely used test of an animal’s ability to retrieve food pellets (Whishaw & Pellis, 1990), we constructed a device to present food pellets on 3D-printed trays atop a conveyor system that sequentially presented pellets in set positions relative to narrow retrieval slits (**Fig. 1A**). Pellets were presented in three-walled compartments atop pillars, a geometry that required grasp-and-lift retrieval and prevented dragging solutions (Bowles et al., 2021; Fenrich et al., 2016); pilot experiments showed this geometry to detect injury deficits more reliably than flat or recessed surfaces (**Supp. Fig. 1 and 2**). Visual inspection of post-run trays scored each pellet as missed (still atop pillar), displaced (present but moved from pillar), or retrieved (no longer visible) (**Fig. 1B**). Using customized tray loaders and by placing multiple retrieval cages before each conveyor, a single user could readily administer the task to sixteen animals simultaneously.

**Figure 1.**
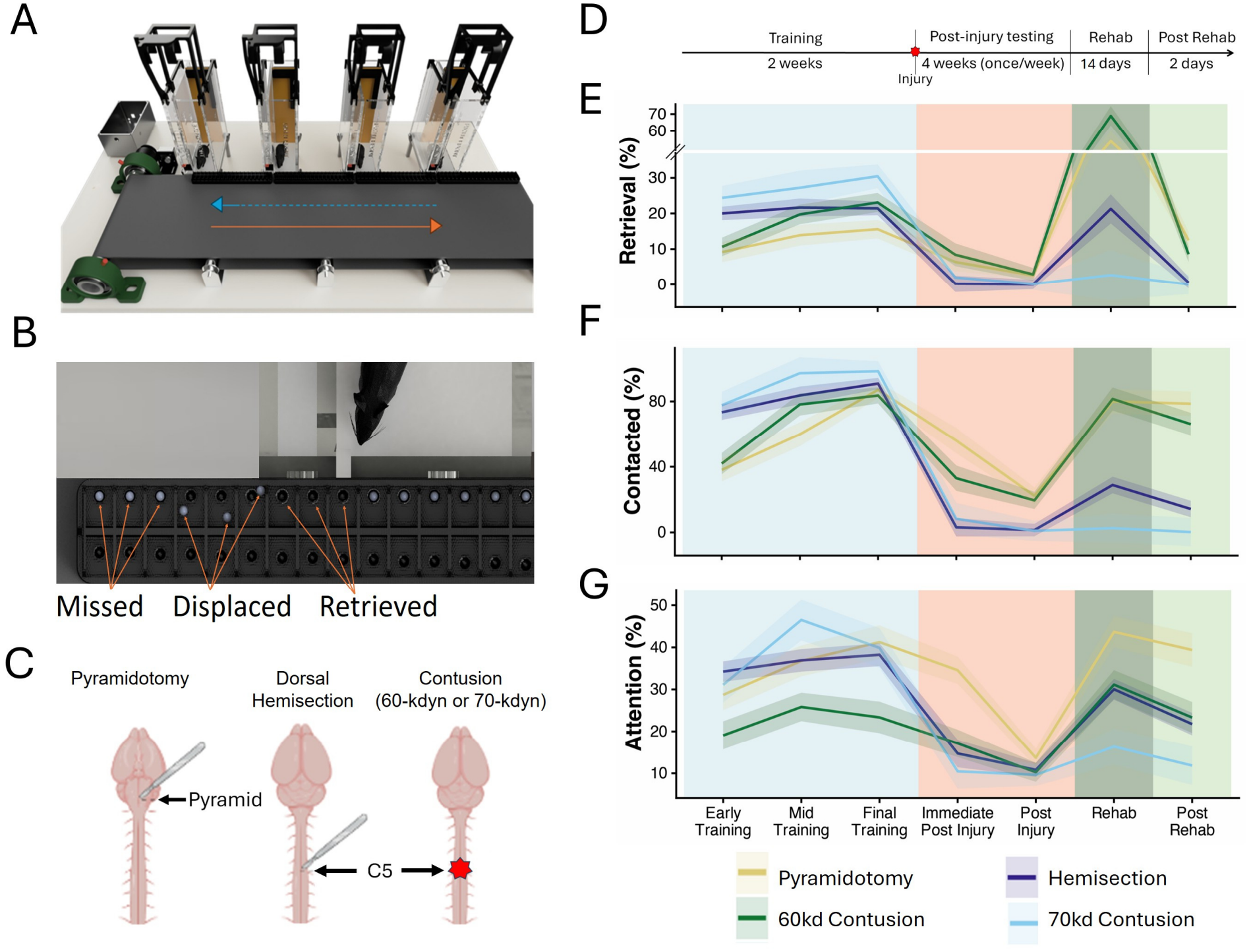
Semi-automated administration of the single pellet retrieval task detects injury-specific forelimb deficits and responsiveness to rehabilitation. (A) Illustration of multiple retrieval boxes, each with camera mounts, interfaced with a programmable conveyor for pellet presentation. (B) Pellets are presented sequentially to mice on 3D-printed trays and then scored as missed (still atop pillar), displaced (off pillar but in compartment) or retrieved (absent). (C) Illustration of injury types (D) Experimental timeline (E-G) Average values of (E) percent of pellets retrieved (F) percent of pellets contacted and (G) percent of time attending to the task in each of four injury types as they progress through experimental phases. Line shading represents SEM. Table 3 provides all pairwise statistical tests. N=12 (pyramidotomy), 33 (transection), 14 (60-kdyn contusion) and 9 (70kdyn contusion).

We first used this standard platform to directly compare deficits in retrieval caused by four different injury types: pyramidotomy, cervical dorsal hemisection, and contusion of 60kdyn or 70kdyn impact force. We also compared the degree to which deficits could be rescued by a paradigm of task-specific rehabilitation (**Fig. 1C**). Each of the four groups moved through a standardized sequence of experimental phases: pre-training, injury, post-injury testing, intensive rehabilitation, and final post-rehabilitation testing (**Fig. 1D**). Injuries were verified by PKCγ for the pyramidotomy group (**Supp. Fig. 3**), GFAP IHC in the hemisection group (**Supp. Fig. 4**), and by volumetric assessment of cleared tissue in contusion groups (**Supp. Fig. 5**). Differences in percent pellet retrieval and contact (retrieved + displaced) were tested by mixed linear models in which injury, experiment phase, and their interaction were set as fixed effects, followed by post-hoc pairwise comparisons. Before injury the four groups performed similarly, with indistinguishable rates of pellet contact (all pairwise comparison p>.05) and overall concordance in retrieval, although the 70-kdyn group showed higher pre-injury success. The cohorts therefore entered the study with closely matched performance, establishing a common baseline against which injury and recovery could be measured. All four types of injury significantly reduced performance (p<.05, pre-injury vs. post-injury), but to varying degrees (**Fig. 1E, F**). Pyramidotomy and 60-kdyn groups were less affected by injury, showing approximately 85% and 75% reductions in retrieval and contact success, respectively, while dorsal hemisection and 70-kdyn groups were more strongly affected, with both retrieval and contact rates reduced to near zero (p<.05 for pairwise comparisons between post-injury pyramidotomy or 60kdyn contusion versus hemisection or 70kdyn). See **Table 2** for all session values and **Table 3** for the complete set of pair-wise tests. Thus, when directly compared in the same assay, dorsal hemisection injuries and 70-kdyn contusion injuries caused more pronounced deficits in forelimb function than pyramidotomy or 60-kdyn contusion injuries.

We next asked whether rehabilitation in the form of intensive daily practice could improve task performance, and whether responsiveness to rehab differed between injury types. Importantly, rehabilitation was performed at least five weeks post-injury and at a time when injury deficits were stable (**Supp. Fig. 6**; no change across prior three sessions, type III ANOVA on the larger linear mixed model following Satterthwaite’s method, main effect of time, retrieval, F(2, 70.2) = 0.97, p = 0.383; contact, F(2, 52.4) = 2.20, p = 0.121), thus providing a consistent baseline to detect potential rehabilitation effects. Rehabilitation was highly effective in pyramidotomy injuries, restoring both contact and retrieval rates to pre-injury levels (p>.05, post-rehab vs. pre-injury). At the other extreme, animals with severe 70-kdyn injuries showed no significant recovery after rehabilitation (p>.05, post-rehab vs. post-injury) with both contact and retrieval rates remaining near zero. The effects were intermediate in the other two injury types. In 60-kdyn animals, rehabilitation restored contact rates nearly to pre-injury levels but produced only partial gains in retrieval, with final values differing significantly from both pre-rehab and their initial baseline (p<.05). Finally, in dorsal hemisection animals, rehabilitation produced a significant but partial improvement in pellet contact (p=.013) but no improvement in pellet retrieval (p=.805, post-rehab vs. post-injury). Post-hoc comparisons confirmed significant differences in the size of rehabilitation effects between different injury types (**Table 3**). In summary, these results demonstrate large differences in the effects of intensive rehabilitation on forelimb function across four injuries of graded severities, ranging from no improvement to complete restoration of task success.

### Automated tracking of reach kinematics

To examine how injury and rehabilitation affect reaching movements, we recorded every session from above and built an analysis pipeline that tracks body parts and pellet location (**Fig. 2A**), identifies individual reaches, and extracts their kinematics (**Fig. 2B**). Tracking the nose also allowed continuous monitoring of task engagement (attention score), and tracking the pellet allowed each reach to be assigned an outcome (miss, displacement, or retrieval). Across all sessions, the pipeline identified 159,343 reaches.

**Figure 2.**
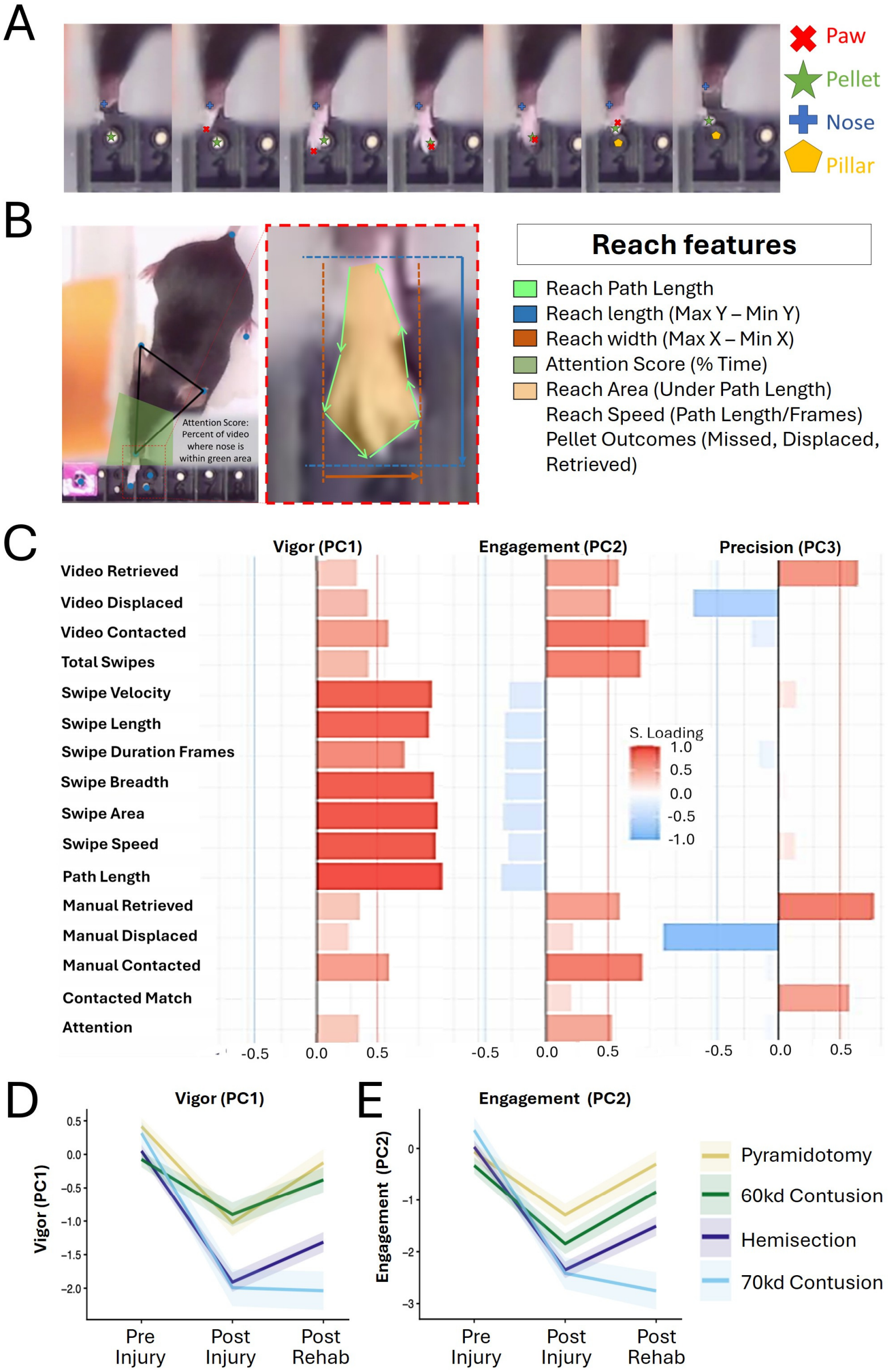
Automated motion tracking and principal component analysis reveal clusters of interrelated variables with injury-specific responsiveness. (A) Top-down recording enables continual detection and tracking of nose, paw, pillar, and pellet during reaching events. (B) Illustration of attention (% of time the nose is positioned in the retrieval slit) and representative kinematic features. (C) shows the loading of individual variables onto the first three principal components. Red and blue indicate positive and negative loadings, respectively. (D,E) show changes in Vigor (D) and Engagement (E) in four different injury groups as mice progress through main experimental phases. Lines represent groups averages and shading represent SEM. Pairwise statistical comparison are provided in Table 6.

### Task Engagement

Changes in task performance after injury and rehabilitation could reflect changes in motor ability and/or changes in motivation due to shifts in effort required per reward. We therefore monitored task engagement based on the percentage of each session the animal spent positioned at the slit (attention score, **Fig. 1G**). Like retrieval and contact, attention showed significant main effects of injury and phase and an injury-by-time interaction (linear mixed model: F(6, 56.6) = 3.98, adjusted p = 0.002). Injury significantly reduced attention in all groups (p<.05, pre-injury vs. post-injury), indicating that animals spent less time attempting the task when injury made retrieval more difficult. Interestingly, although suppressed from baseline, attention was higher in the first post-injury testing session compared to subsequent weeks (**Fig. 1G**), consistent with task extinction as reward diminished. Overall post-injury attention was similar between groups (p > 0.32, all group comparisons in the post-injury phase), and after the drop observed following the first test session was stable in the weeks preceding rehabilitation (**Supp. Fig. 6**; main effect of time, F(2, 50.7) = 0.08, p = 0.919). Rehabilitation strongly affected attention in some groups, again with effects that paralleled task outcomes. In pyramidotomy and 60-kdyn groups, rehabilitation significantly increased attention until it no longer differed from pre-injury baseline (p>0.728, post-rehab vs. baseline). After dorsal hemisection attention improved partially, remaining significantly different from both baseline and post-injury (p<.001). Finally, rehabilitation produced no change in the 70-kdyn group (p=0.697, pre-rehab vs. post-rehab) matching the lack of improvement in task performance. Thus, in addition to affecting overall task success, injury and rehabilitation influence the amount of time that animals choose to spend attempting the task, with effects that depend on the type of injury.

### Principal component analysis of reaching behavior

We next applied nonlinear principal component analysis (PCA) to reach-level data to identify relationships between sixteen variables covering of kinematic qualities, attention, digitally scored reach outcomes, and manually scored session retrieval and contact (see Methods). Outcome measures were included to capture potential relationships between kinematics, engagement, and task success. The Kaiser criterion, scree plot, and permutation testing (1,000 permutations) all supported a three-component solution that accounted for 64% of the total variance (**Fig. 2C**). PC1 (32% of the total variance) loaded heavily on the kinematic features involving the speed and extent of the reach and somewhat on pellet contact, but not retrieval; we termed it “Vigor.” PC2 (18%) loaded strongly on attention, total reaches, and pellet contact and was termed “Engagement.” PC3 (14%) loaded positively on retrieval and negatively on displacement and was therefore termed “Precision.”

We then tested whether Vigor, Engagement, and Precision scores differed by injury group and phase. For Vigor and Engagement, RM-ANOVA identified significant main effects of injury, phase, and their interaction. All injuries significantly reduced both Vigor and Engagement (p < .05, baseline vs. post-injury; Holm-adjusted estimated marginal means). As with task outcomes, the injuries sorted into two tiers. Vigor and Engagement were less affected in pyramidotomy or 60-kdyn contusion than in dorsal hemisection or 70-kdyn contusion groups, as shown by significant differences in pairwise comparison made between injuries between but not within tiers (p<.05). Interestingly, this post-injury grouping seen in Engagement differed from the similarity in attention scores (above), likely reflecting the incorporation of additional variables like total reaches into the Engagement PC2. Rehabilitation improved Vigor and Engagement in pyramidotomy and 60-kdyn animals to near baseline values, and in the dorsal hemisection group produced a significant but partial improvement. In contrast, rehabilitation produced no improvement in Vigor or Engagement in 70-kdyn animals. Regarding Precision (PC3), main effects were detected only for injury (F(3, 50) = 14.56, p < 0.001, η^2^ = 0.47) and experimental phase (F(2, 100) = 11.06, p < 0.001, η^2^ = 0.18) but not their interaction (F(6, 100) = 1.84, p = 0.110, η^2^ = 0.10), and were driven largely by a difference between pyramidotomy and the other three groups and by a difference between post-injury versus the other two phases. Thus, unlike Vigor and Engagement, Precision does not capture type-specific responses to injury or rehabilitation. Overall, these results separate the effects of injury and rehabilitation on movement vigor from their effects on task engagement, with both depending on the type and severity of injury.

### Kinematic features that predict retrieval

To complement the PCA we next identified specific kinematic features that were most closely associated with task success. Using reaches from the pre-injury baseline phase, we used least absolute shrinkage and selection operator (LASSO) regression with 10-fold cross-validation to identify features of reaching (path over frames [speed of reach], swipe length, path length, speed, area, breadth, total swipes, and swipe duration) that most strongly predicted the outcome (retrieval, displacement, or miss) in animals prior to injury. At λ = 0.020, six of the eight candidate predictors retained non-zero coefficients, with swipe speed and swipe duration reduced to zero and therefore excluded by the LASSO. The selected model had R^2^ = .134 and a 10-fold cross-validated mean squared prediction error of .866 (SE = .004), compared with R^2^ = .135 and prediction error = .865 for the unregularized seven-predictor model. Three features emerged as most predictive: length, area, and speed of reach (**Fig. 3A, B**), with longer, larger, and faster reaches associated with retrieval. Because all three features also load on Vigor, they can be viewed as concrete, measurable components of that composite.

**Figure 3.**
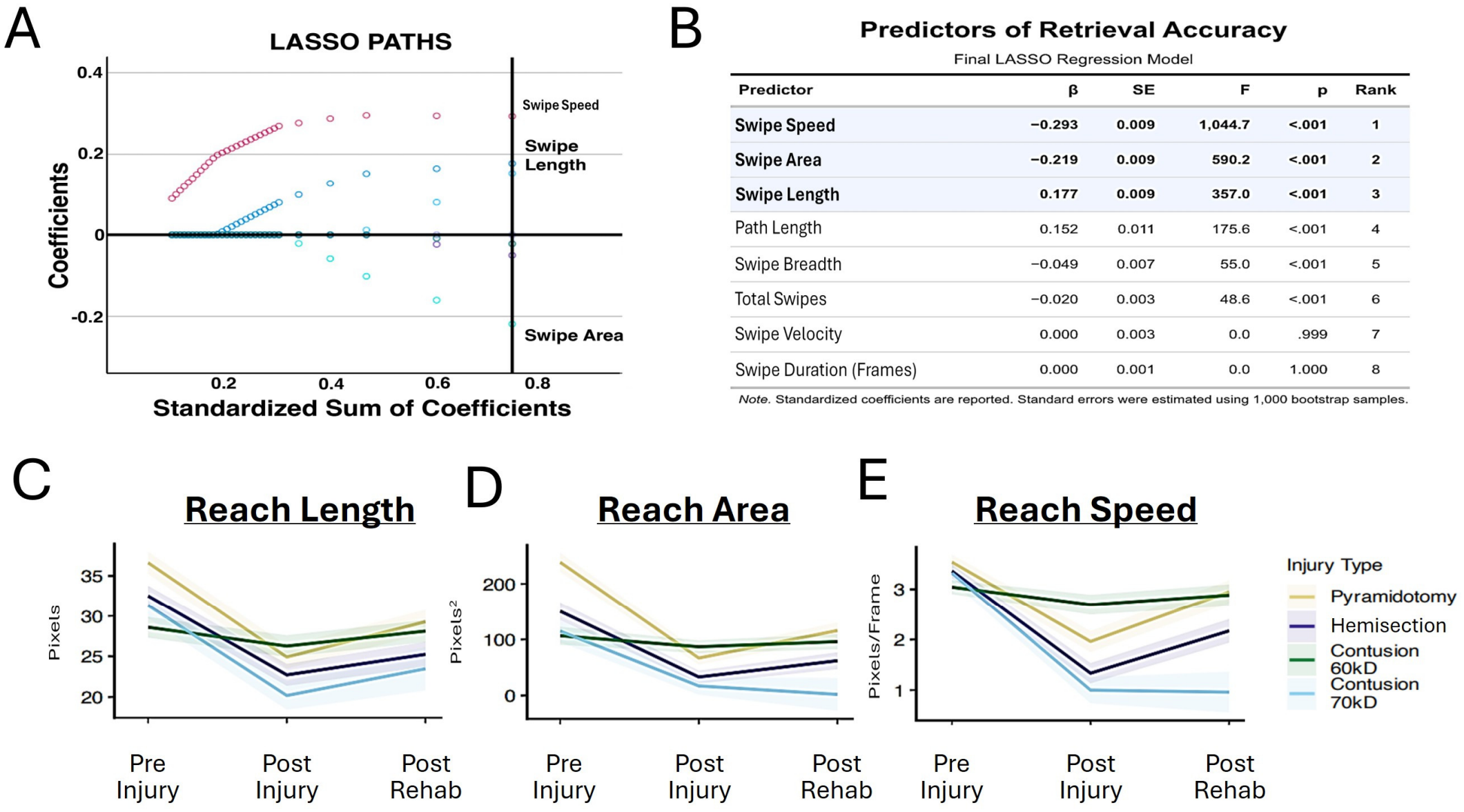
LASSO selection of predictors of retrieval accuracy. (A) LASSO coefficient paths. Lines indicate the standardized coefficient of each predictor as the penalty is relaxed, plotted against the standardized sum of coefficients. The vertical line marks the optimal model, selected by 10-fold cross-validation. The three predictors with the largest retained coefficients are labeled (swipe length, swipe area, swipe speed). (B) Final LASSO regression model at the optimal penalty, with predictors ranked by the magnitude of their effect. Shaded rows are the three strongest predictors (swipe speed, swipe length, swipe area). Swipe velocity and swipe duration (frames) were shrunk to zero. (C-E) show changes in lenth (C), Area (D), and speed (E) in four different injury groups as mice progress through main experimental phases. Lines represent groups averages and shading represent SEM. Pairwise statistical comparisons are provided in Table 3.

We then tested how each was affected by injury and rehabilitation. All three features showed significant injury-by-phase interaction (linear mixed models: length, F(6, 52.5) = 3.71, adjusted p = 0.004; area, F(6, 50.5) = 6.10, adjusted p < 0.001; speed, F(6, 53.8) = 7.72, adjusted p < 0.001), indicating that their response to injury depended on injury type. In pyramidotomy animals, all three features were significantly reduced by injury (p<.01, baseline vs. post-injury) and significantly elevated by rehabilitation (p<.01, pre-vs. post-rehabilitation), tracking the improvement in task performance in this group. In dorsal hemisection animals all three features were suppressed by injury (p<.01, baseline vs. post-injury), but only reach speed improved after rehabilitation (speed p<.001, area p=.06, length p=0.174, pre-rehab vs. post-rehab). This recovery in speed paralleled the partial recovery in pellet contact, even though retrieval did not improve. After 70-kdyn contusion all three features declined (p<.001, baseline vs. post-injury) and none improved with rehabilitation (p>0.284), consistent with the persistent deficits in task outcomes. In contrast, the 60-kdyn group showed no overall change in average reach length, area, or speed after injury or rehabilitation (**Fig. 3C, D, E**). Closer examination, however, revealed that all three features were highly dynamic in nearly all 60-kdyn animals across the experimental phase but with bidirectional effects (**Supp. Fig. 7A, B, C**). For example, while eight of fourteen animals showed a reduction in reach length of at least 10% after injury, four others showed an increase larger than 10%, resulting in a non-significant change in group average (p=0.488, baseline vs post-injury) (**Supp. Fig. 7A, B, C**). This magnitude of change exceeded the variability in pre-injury sessions. These results indicate a complex relationship between moderate contusion injury and success-predictive kinematic features and highlight the importance of monitoring individual variability in kinematic features. Overall, for most injury types, LASSO regression identified specific features of motion that correlate with the success of pellet retrieval prior to injury, which are suppressed by damage and show variable response to rehabilitation across injury type.

## DISCUSSION

We present a semi-automated device for rapid, multi-animal administration of the single pellet retrieval task (SPRT) with integrated extraction of kinematic measurements from thousands of individual reach events and an associated PCA/LASSO analysis pipeline to discover relationships among these measurements and the reach features most closely tied to task success. The design was chosen to balance cost, simplicity, and animal throughput against data richness. It offers an alternative to systems that provide more detailed reach monitoring at limited animal throughput (Berezhnoi et al., 2024; Bowles et al., 2021; J. Z. Guo et al., 2015; Palmér et al., 2012) and to single-unit robotic automation systems with higher device complexity and limited kinematic readout (Kakanos et al., 2023; Torres-Espín et al., 2018). The result was a dataset of kinematic features extracted from more than 150,000 reaches from 62 animals across four injury types.

This throughput allowed injuries to be measured on the same platform, enabling direct comparison of kinematic and success outcomes. Interestingly, the same rehabilitation paradigm restored full retrieval success in one injury (pyramidotomy), showed no effect in another (70-kdyn contusion), and intermediate effects in other injuries, thus defining conditions in which the form of training is effective. The results also illustrate the sensitivity of kinematic profiling compared to outcome-only readouts. For example, in the pyramidotomy group key kinematic features showed significant but partial improvements after rehabilitation whereas retrieval success itself recovered fully. Conversely, in the dorsal hemisection group some kinematic features showed partial restoration toward baseline while retrieval success was unchanged. In line with prior work, these results support the ability of the present kinematic analyses to detect both deficits and partial improvements that are not apparent in the net task outcome (Lai et al., 2015; Whishaw, 2000).

A main finding of the PCA was a separation between motion characteristics and task engagement. PC1 (Vigor) was influenced mostly by the characteristics of motion, with strong positive loadings for speed and size of reach and a moderate loading for pellet contact, and was therefore termed Vigor. In contrast, PC2 reflected the amount of time the animal was positioned to perform the task, the number of reaches performed, and also loaded strongly for pellet contact regardless of retrieval outcome. Injury reduced both PCs, indicating that injury affects not only the motor ability to execute retrieval but also the willingness of the animal to attempt the task, possibly reflecting a shift in effort-to-reward ratio. This interpretation is strengthened by direct measurements of post-injury attention, which fell sharply after injury but was partly maintained in the testing session immediately after injury, when animals are first adjusting to injury-induced difficulty in retrieval. Attention scores fell more in subsequent post-injury tests, perhaps reflecting a decrease in expected reward. The rehabilitation paradigm, which began with easier tray configurations and progressed back to pillars, also increased both components, indicating that it influenced motivation as well as motor performance. PC3 (Precision), which captured retrieval relative to displacement, declined after injury and partially improved after rehabilitation, but without significant differences between injury types. Overall, these results highlight the importance of measuring motor and engagement effects separately when evaluating injury and treatment.

PCA is valuable for revealing relationships within data, but its composite outputs are abstract and do not always lead to mechanistic understanding or specific targets for optimization (Jollife & Cadima, 2016; Ma & Dai, 2011; Reinhardt et al., 2010). One common solution is to manually select individual variables as concrete surrogates for each component; here, for example, attention may stand in for PC2 (Engagement). A more general solution is a data-driven method that ranks individual variables by their relationship to the outcome of interest. We used LASSO for this purpose, identifying specific kinematic features most strongly associated with outcomes of reaching events. This approach indicates that retrieval is most likely when reaches are rapid and reach full extension. These features declined after injury, and their recovery tracked task recovery after pyramidotomy and, in part, after dorsal hemisection. This latter observation addresses a central question in neuro-recovery, whether an intervention improves task outcomes by restoring the original movement (“true recovery”) or by promoting alternative, compensatory motions (Almli & Finger, 1988; Lai et al., 2015; Torres-Espín et al., 2018; Whishaw, 2000). By identifying key kinematic features most strongly associated with task success, and then monitoring these features after injury and rehabilitation, the LASSO approach indicates that in some injuries, particularly pyramidotomy and dorsal hemisection, the present rehabilitation paradigm acts at least in part through the restoration of high-value motion characteristics.

This study has several limitations. Kinematics were measured from a single top-down camera, so they capture the two-dimensional trajectory of the paw but not digit movements, grasp, or vertical motion. The attention score is a proxy for motivation and cannot separate reduced willingness from reduced ability, pain, or fatigue. We tested a single rehabilitation paradigm, so the boundaries of efficacy described here may differ with other training regimens. Finally, lesion volume was measured in a subset of contusion animals and was not related to behavior at the level of individual animals.

In summary, this work presents a simple, flexible platform for high-throughput administration of the single pellet retrieval task, with automated kinematic extraction that scales to more than 150,000 reaches. The accompanying PCA/LASSO framework reveals how behavioral measures relate to one another and identifies the specific variables most relevant to task success. Together, these hardware, software, and analysis tools address the twin challenges of collecting and interpreting complex behavioral data after nervous system injury.

