## Supplemental Figures for "Automated, high-throughput kinematic analysis of skilled reaching in rodent models of neurological injury"

### Supplemental Figure 1

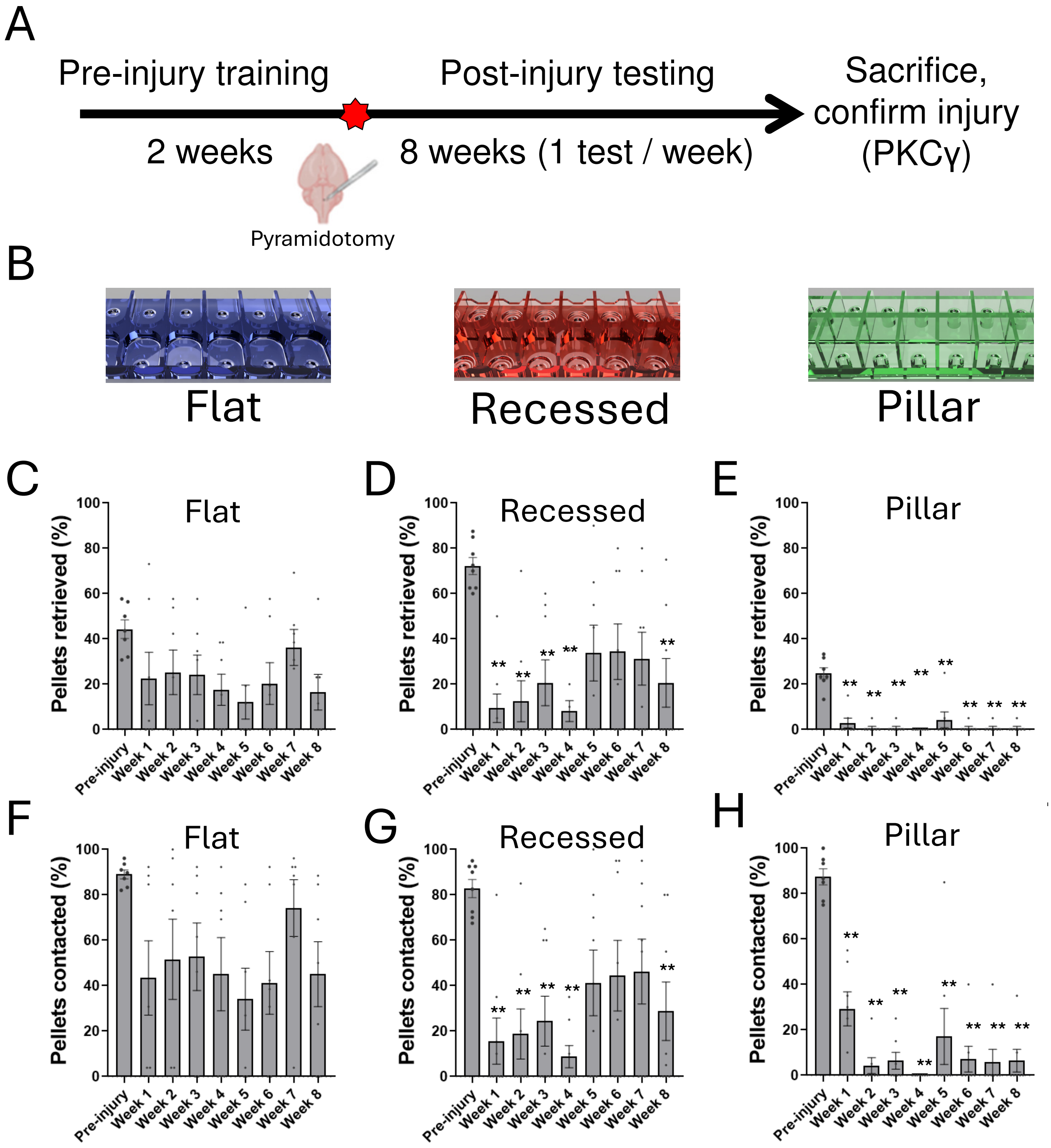


**Supplemental Figure 1. Tray geometry affects forelimb reaching outcomes.** (A) Experimental timeline (B) illustrations of three tray geometries in which pellets are presented on flat surfaces (blue), in recessed holes (red), or atop pillars (green). In all cases each pellet is presented in a separate three-walled compartment. (C-E) show pellet retrieval as a percent of total pellets during pre-injury testing (average of final three tests) and in weekly tests performed for eight weeks post-injury. (F-H) show the percent of pellets that were contacted prior to injury and each week post-injury. Only pillar trays shows strong and significant reduction in both pellet retrieval and pellet contact at all time points tested. **p<.01, one-way ANOVA with post-hoc Tukey's. N=7 (flat), 8 (recessed), 7 (pillar).

### Supplemental Figure 2

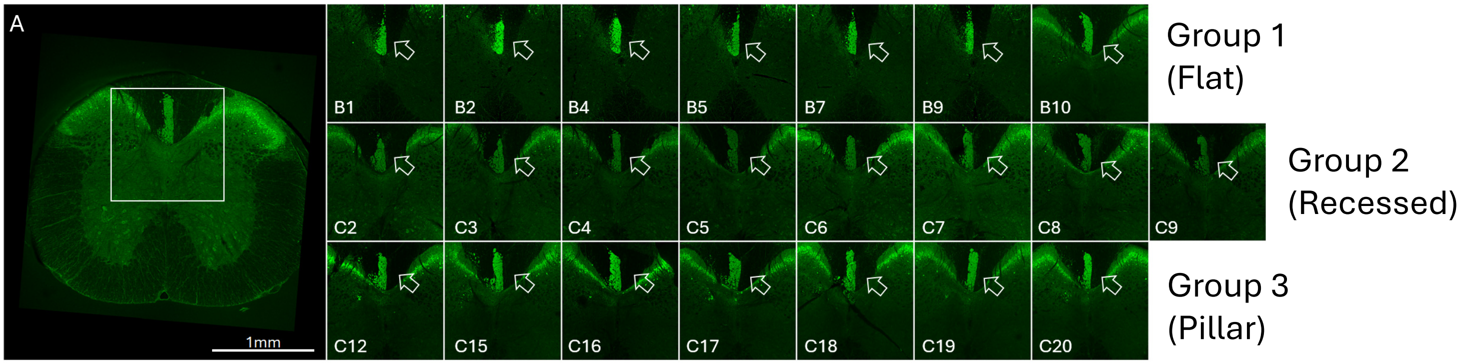


**Supplemental Figure 2. PKC-gamma immunohistochemistry in cervical spinal cord confirms unilateral ablation of the corticospinal tract in animals used for tray optimization.** (A) shows transverse section of cervical spinal cord with detection of PKC-gamma, a protein expressed in corticospinal tract axons, in green. The white box indicates the imaging area in subsequent panels. (B1-C20) Images of dorsal columns show the left corticospinal tract labeled by PKC-gamma (green) and undetectable signal on the right (open arrows), confirming unilateral CST loss in all animals. Panel labels correspond to animal ID.

### Supplemental Figure 3

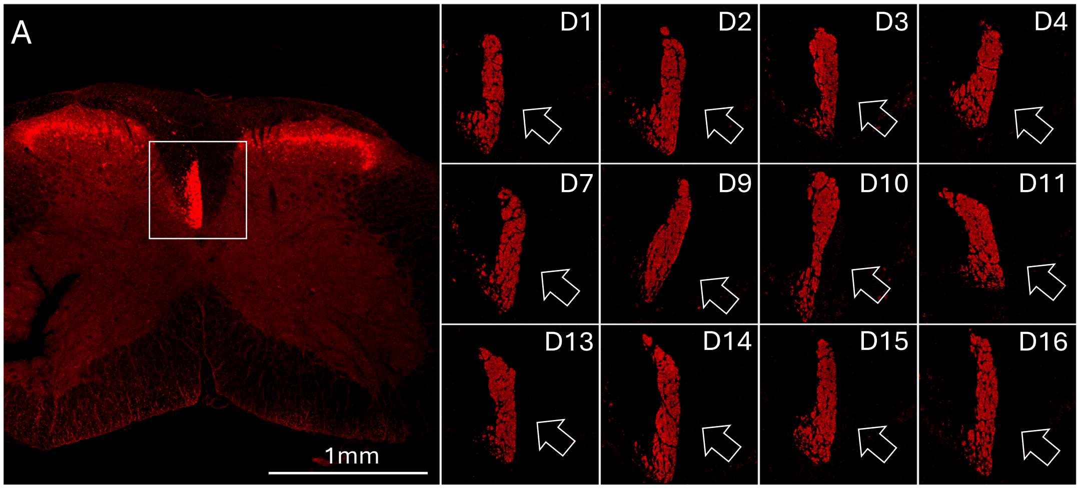


**Supplemental Figure 3. PKC-gamma immunohistochemistry in cervical spinal cord confirms unilateral ablation of the corticospinal tract.** (A) Transverse section of cervical spinal cord with detection of PKC-gamma, a protein expressed in corticospinal tract axons, in red. The white box indicates the imaging area in subsequent panels. (D1-D16) Images of dorsal columns show the left corticospinal tract labeled by PKC-gamma (red) and undetectable signal on the right (open arrows), confirming unilateral CST loss in all animals. Panel labels correspond to animal ID.

### Supplemental Figure 4

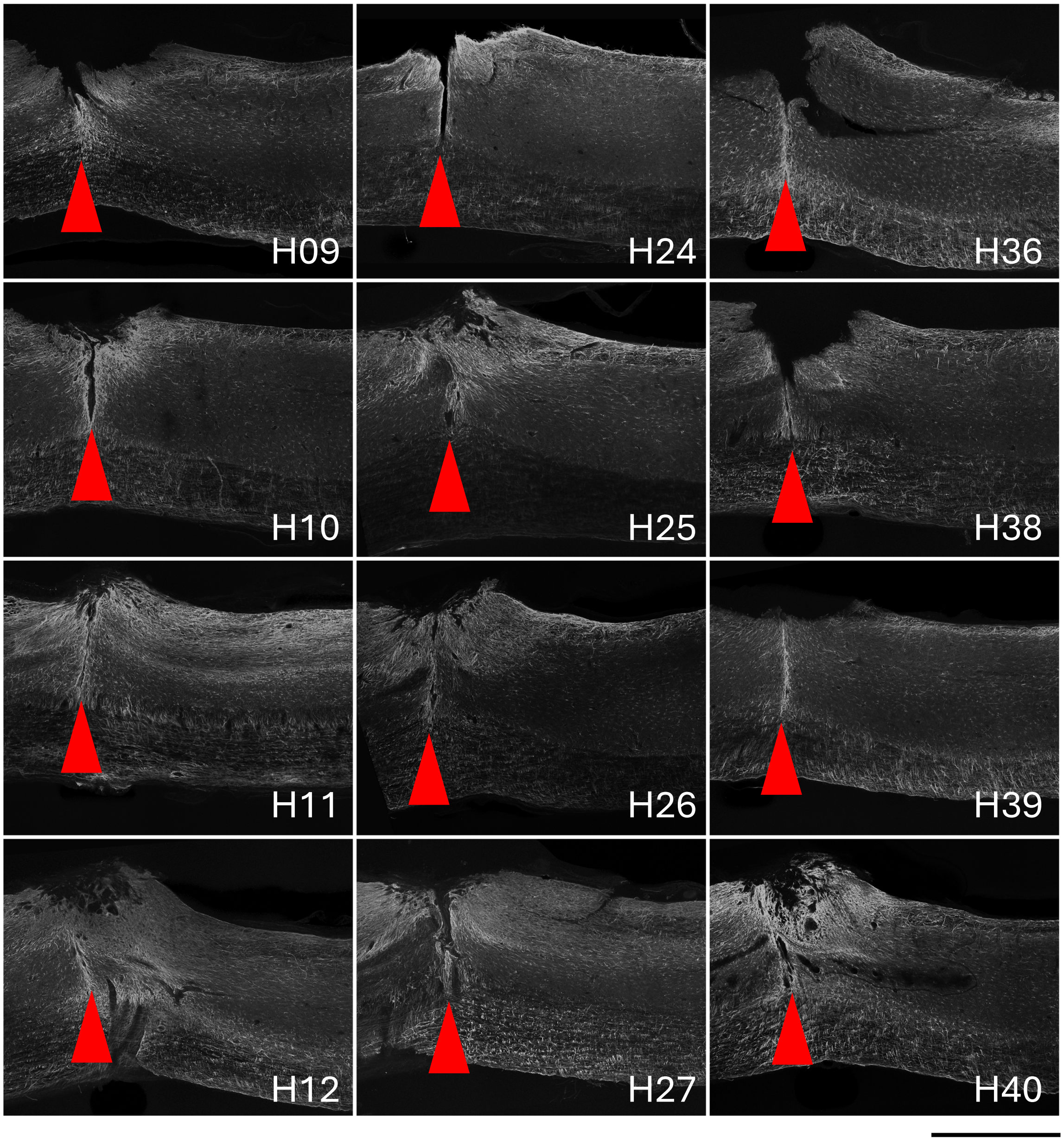


**Supplemental Figure 4. GFAP immunohistochemistry in cervical spinal cord confirms transection injury.** Images are sagittal sections of cervical spinal cord with reactive gliosis detected by immunohistochemistry for GFAP. Red arrows indicate detection of spinal transection in each animal. Panel labels correspond to animal ID. Scale bar is 1mm.

### Supplemental Figure 5

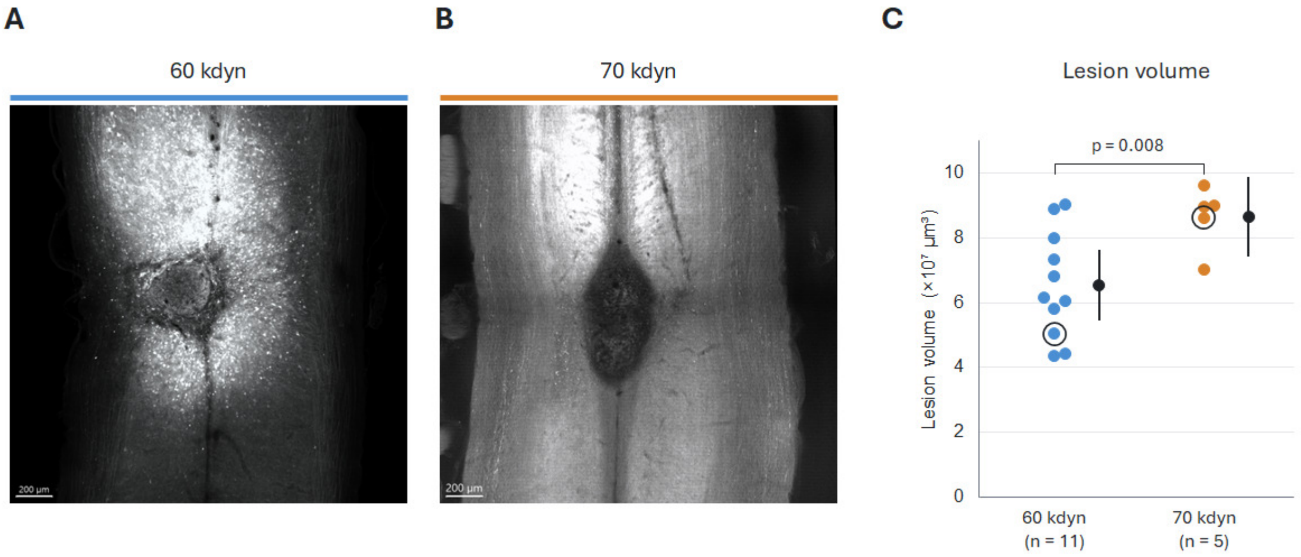


**Supplemental Figure 5. Lesion volume after 60 and 70 kdyn spinal cord contusion.** (A, B) Representative Imaris renderings of tissue autofluorescence (561 nm excitation) at the injury epicenter after a 60 kdyn (A) or 70 kdyn (B) contusion (spinal level, time post-injury]) The lesion is the region of altered autofluorescence intensity and irregular texture at the center of each image. Scale bars, 200 µm. (C) Lesion volume measured in Imaris from manually traced 3D surfaces. Each point is one animal; black points and bars show the mean ± 95% confidence interval. Circled points are the animals shown in A and B. 60 kdyn, n = 11; 70 kdyn, n = 5. Welch's two-sided t-test, p = 0.008.

### Supplemental Figure 6

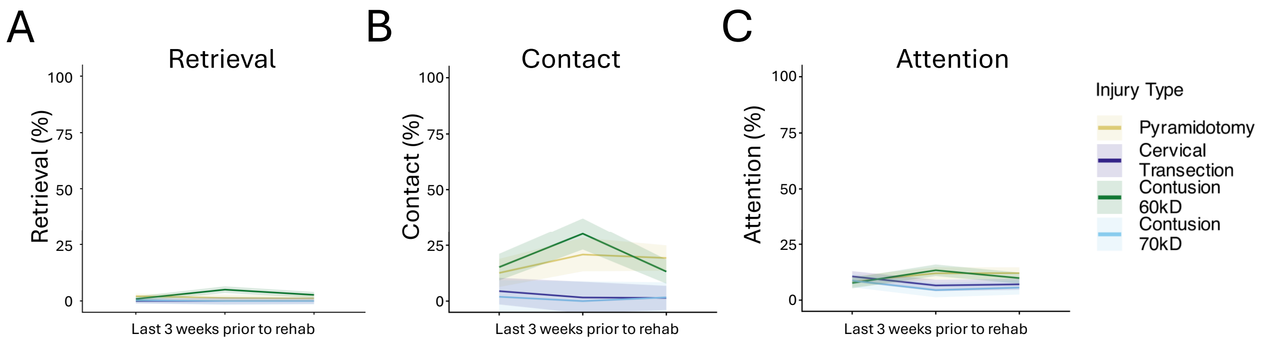


D

Retrieval

```
> anova(mod_retrieved, type = 3)
Type III Analysis of Variance Table with Satterthwaite's method
                                   Sum Sq Mean Sq NumDF  DenDF F value Pr(>F)
FINAL_THREE_INJURYWEEKS            24.492  12.246     2 70.234  0.9728 0.3831
Injury_type                        66.055  22.018     3 48.689  1.7491 0.1693
FINAL_THREE_INJURYWEEKS:Injury_type 110.716 18.453    6 67.991  1.4659 0.2031
```

E

Contact

```
> anova(mod_contacted, type = 3)
Type III Analysis of Variance Table with Satterthwaite's method
                                   Sum Sq Mean Sq NumDF  DenDF F value Pr(>F)
FINAL_THREE_INJURYWEEKS            386.88  193.44     2 52.354  2.1962 0.1214
Injury_type                        816.85  272.28     3 49.308  3.0913 0.0354 *
FINAL_THREE_INJURYWEEKS:Injury_type 1209.33 201.56    6 58.117  2.2883 0.0474 *
```

F

Attention

```
> anova(mod_attention, type = 3)
Type III Analysis of Variance Table with Satterthwaite's method
                                   Sum Sq Mean Sq NumDF  DenDF F value  Pr(>F)
FINAL_THREE_INJURYWEEKS              3.62   1.811     2 50.698  0.0841 0.91942
Injury_type                         58.16  19.385     3 49.264  0.9007 0.44761
FINAL_THREE_INJURYWEEKS:Injury_type 361.07  60.178    6 50.331  2.7959 0.02001 *
---
```

**Supplemental Figure 6. Performance and attention in the pellet retrieval task are stable prior to rehabilitation.** (A-C) show average pellet retrieval (A), pellet contact (B), and task attention (C) plotted for three weekly testing sessions performed prior to initiation of rehabilitation. (D-F) Statistical analysis identifies no significant difference across time in any group. Thus, any subsequent change after rehabilitation reflects the effect of intervention and not a continuation of any pre-existing trajectory of task improvement.

### Supplemental Figure 7

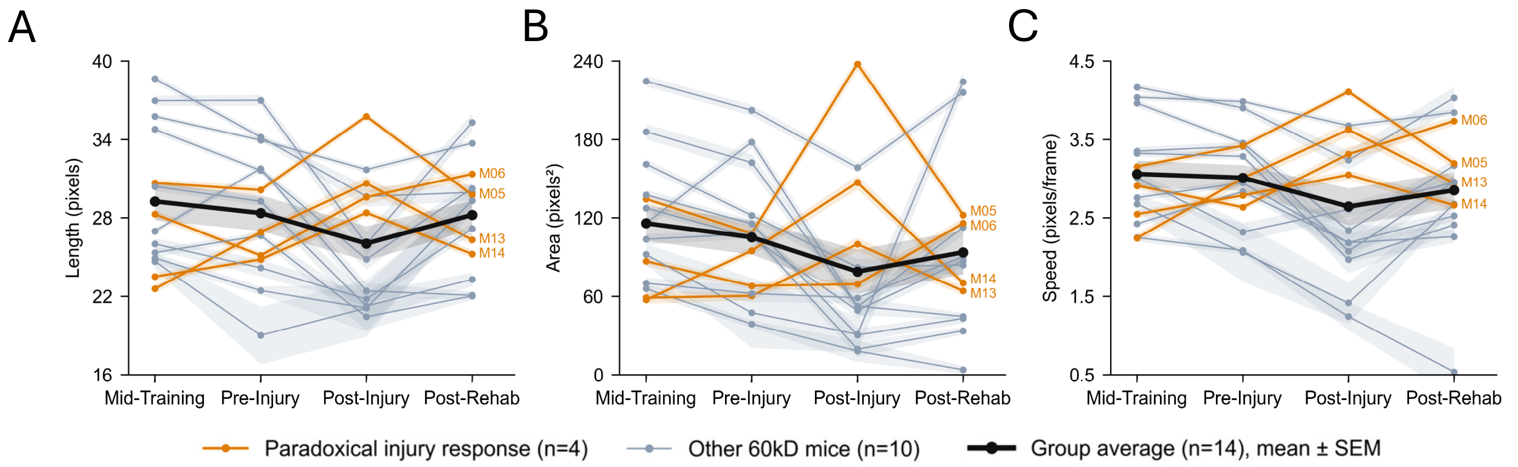


**Supplemental Figure 7. 60-kdyn contusion injuries and rehabilitation affect key kinematic features of reach with variable directionality in individual animals.** Each panel shows average values for individual animals across main experimental phases. (A) shows reach length, (B) shows reach area, and (C) shows reach speed. In most animals these features were stable prior to injury (compare mid training to pre-injury) and altered after injury and again after rehabilitation. Four animals, colored orange, showed an unusual increase in these features after injury and decrease after rehabilitation, the opposite pattern from most other animals in the rest of the cohort and in other injury groups. Thus, the group average (black line) shows little change between experimental phases.
